# Including intraspecific trait variability to avoid distortion of functional diversity and ecological inference: lessons from natural assemblages

**DOI:** 10.1101/2020.09.17.302349

**Authors:** Mark K. L. Wong, Carlos P. Carmona

## Abstract

1. Functional diversity assessments are crucial and increasingly used for understanding ecological processes and managing ecosystems. The functional diversity of a community is assessed by sampling traits at one or more scales (individuals, populations, species) and calculating a summary index of the variation in trait values. However, it remains unclear how the scale at which traits are sampled and the indices used to estimate functional diversity may alter the patterns observed and inferences about ecological processes.
2. For 40 plant and 61 ant communities, we assess functional diversity using six methods – encompassing various mean-based and probabilistic methods – chosen to reflect common scenarios where different levels of detail are available in trait data. We test whether including trait variability at different scales (from individuals to species) alter functional diversity values calculated using volume-based and dissimilarity-based indices, Functional Richness (FRic) and Rao, respectively. We further test whether such effects alter the functional diversity patterns observed across communities and their relationships with environmental drivers such as abiotic gradients and occurrences of invasive species.
3. Intraspecific trait variability strongly determined FRic and Rao. Methods using only species’ mean trait values to calculate FRic (convex hulls) and Rao (Gower-based dissimilarity) distorted the patterns observed when intraspecific trait variability was considered. These distortions generated Type I and Type II errors for the effects of environmental factors structuring the plant and ant communities.
4. The high sensitivity of FRic to individuals with extreme trait values was revealed in comparisons of different probabilistic methods including among-individual and among-population trait variability in functional diversity. By contrast, values and ecological patterns in Rao were consistent among methods including different scales of intraspecific trait variability.
5. Decisions about where traits are sampled and how trait variability is included in functional diversity can drastically change the patterns observed and conclusions about ecological processes. We recommend sampling the traits of multiple individuals per species and capturing their intraspecific trait variability using probabilistic methods. We discuss how intraspecific trait variability can be reasonably estimated and included in functional diversity in the common circumstance where only limited trait data are available.

## INTRODUCTION

Assessments of the diversity of organisms’ functional traits – ‘functional diversity’ – are important for understanding manifold phenomena ranging from macroevolutionary processes (Diaz et al., 2016; Pigot et al., 2020) to community assembly (McGill et al., 2006) and biodiversity-ecosystem function relationships (Gross et al., 2017). Most functional diversity assessments at and above the community level use just a single value for each trait of each species – the species-mean (Villéger et al. 2008, Mouchet et al. 2010). Calculating functional diversity from species-mean trait values lightens demands on trait measurement, especially when diverse ecological communities and large spatial scales are involved. In studies where trait data for species vary in origin or structure (e.g., Díaz et al., 2016; Pigot et al., 2020), the species-mean trait value can also be used directly to achieve uniform representation across species, facilitating interspecific comparisons.

The species-mean trait value, however, overlooks trait variability among conspecific individuals, which may be extensive due to effects from local adaptation, phenotypic plasticity, developmental conditions and ontogeny (Des Roches et al., 2018). This intraspecific trait variability can determine species’ ecological interactions (Des Roches et al., 2018; Carmona et al., 2019a), and contribute substantially to community functional diversity (as shown by Albert et al., 2011; Messier, McGill & Lechowicz, 2010; Siefert et al., 2015). Assessments failing to account for intraspecific trait variability may therefore misestimate the levels of functional diversity in reality. However, the extent to which these effects alter the observed patterns of functional diversity across communities and inferences about ecological processes are less explored in empirical systems.

Intraspecific trait variability mainly occurs at three hierarchical scales (Albert et al., 2011), each varying in its contribution to functional diversity and relevance to different community processes. At the broadest scale is trait variability among separate local populations. This generally increases as species are distributed across heterogeneous environments. Thus, processes such as environmental filtering may be better detected in functional diversity assessments incorporating population-level trait variability than those surveying species-level trait variability only (Gross et al., 2013). At a finer scale, trait variability among individuals within the same populations affects communities through biotic interactions. Simulation-based, field and experimental studies on plant communities show that accounting for such individual-level trait variability can improve the detection of reduced niche overlap (Mason et al., 2011; de Bello et al., 2013) and the prediction of coexistence outcomes between competing species (Carmona et al., 2019a). At the finest scale (and outside the scope of this study), trait variability within an individual may also influence community processes (Westneat, Wright & Dingemanse, 2015). Few empirical studies have compared the effects of individual-to population-level trait variability on functional diversity patterns (but see Messier, McGill & Lechowicz, 2010).

Functional diversity indices summarise the variation in traits at the considered scale. Methods to calculate functional diversity indices can be categorised under two groups. Those in the first group use the trait dissimilarity between species to calculate community functional diversity. Widely used dissimilarity-based indices include Rao’s quadratic entropy (Botta-Dukát, 2005), functional dispersion (FDis; Laliberté & Legendre, 2010), mean pairwise distance (MPD; Weiher et al., 1998) and the FD index of Petchey & Gaston (2002). In these, dissimilarity is often calculated based on Gower’s distance, which generally does not incorporate intraspecific trait variability because it uses only the mean trait value for each species (but see Cianciaruso et al., 2009). Gower-based dissimilarity is also affected by the species pool considered, as this determines the range of trait values used to standardize Gower’s distances (de Bello et al., 2013). As a less context-dependent alternative, one can compute trait dissimilarity based on the overlap between the trait probability density functions (TPD) of different species (Carmona et al., 2016a; 2019b). Unlike Gower-based dissimilarity, overlap-based dissimilarity using TPD includes intraspecific trait variability.

Methods in the second group use the position of entities (i.e., individuals or species) in a multidimensional trait space to characterise the boundaries of a hypervolume encompassing all trait values observed in the community. The various Functional Richness (FRic) indices calculated using convex hulls (Cornwell et al., 2006), n-dimensional hypervolumes (Blonder et al., 2018), or TPD functions (Carmona et al., 2016a; 2019b), are examples of such volume-based indices. Whereas a convex hull is defined by the positions of entities with the most extreme trait values, n-dimensional hypervolume and TPD functions estimate a probabilistic hypervolume in which the frequencies of different trait values are accounted for (Carmona et al., 2016a; 2019b).

The different scales at which traits can be sampled, often with limited resources, and the variety of methods for calculating functional diversity indices make functional diversity assessments logistically challenging to implement (van der Plas et al., 2017). Empiricists thus often have to choose, *a priori*, the scales of trait variability to include (e.g. species-level only, or including population and/or individual levels), the indices used and the methods to calculate them – with the aim of achieving the most unbiased representation of functional diversity patterns. Although dissimilarity- and volume-based functional diversity indices such as Rao and FRic are used widely in empirical studies on functional diversity (Mouchet et al., 2010), there is little information about their sensitivity to different scales of trait variability.

In communities of plants in the Mediterranean region and ants in subtropical Asia, we investigate the extent to which excluding different scales of trait variability alter the observed functional diversity patterns and conclusions about the environmental factors driving community structure. We first calculate the FRic and Rao of communities using trait data of the highest available resolution (the greatest number of replicates in the smallest sampling unit, i.e., the plot). These ‘HighRes’ methods include as many scales of trait variability permitted by the data and use probabilistic distributions (TPD functions) of trait values to calculate indices that should best approximate the functional diversity in reality. The HighRes method for plants includes individual-, population- and species-level trait variability, while that for ants includes individual- and species-level trait variability. We then compare the values of FRic and Rao from HighRes methods to those from other commonly used methods for calculating functional diversity, which include fewer scales of trait variability owing to the lower resolution of the trait data available. Finally, we model the relationships of FRic and Rao against environmental variables, and test whether the patterns captured with HighRes methods are distorted when the other methods are used to calculate functional diversity.

## MATERIALS AND METHODS

### Community and trait data

The plant community and trait dataset, from Carmona et al. (2015), comprises abundance data for 51 plant species in each of 40 plots distributed along a slope (average inclination: 25%) in central Spain subjected to a Mediterranean climate. Soils towards the upper part of the slope were shallow and with low nutrient and water availability, while soils towards the bottom of the slope were deeper and far more productive. For 10 individuals of each species in each plot, data for two traits – plant height and specific leaf area – were collected, producing a trait dataset encompassing 2540 individuals.

The ant community and trait dataset, from Wong et al. (2020), comprises frequency-of-occurrence data for 29 ant species in each of 61 plots in an open tropical grassland in Hong Kong. One species, present in 24 plots, was the Red Imported Fire Ant (*Solenopsis invicta*), an invasive species known impact the structure of ant communities (Gotelli & Arnett, 2000). Data for seven morphological traits (summarised in Table 1 in Wong et al., 2020) were collected for ≥10 individuals (mean=11, max.=20) of every species, producing a trait dataset encompassing 319 individuals. This included data from separate sub-castes of polymorphic species (Wong et al., 2020). As far as possible, the selected individuals of each species were chosen to reflect the range of body sizes encountered across all samples. Digital photographs taken at every plot were used to estimate percentage ground cover via colour thresholding techniques in ImageJ (Abramoff, 2004).

**Table 1.**
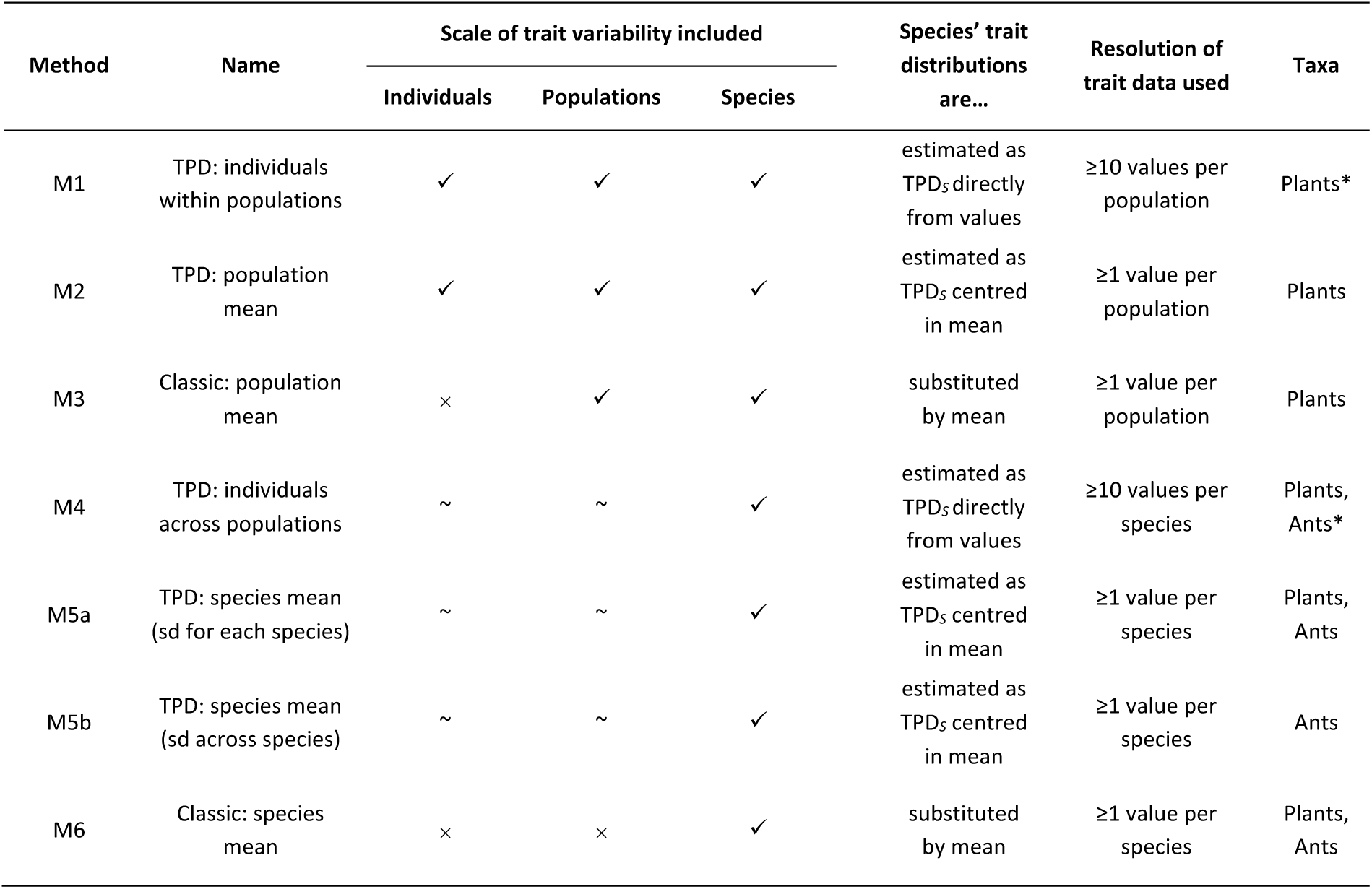
Methods for assessing functional diversity based on the resolution of trait data. A given method may include up to three hierarchical scales of trait variability: among individuals within the same populations (Individuals), among separate local populations (Populations), and among different species (Species). Methods differ in whether trait variability at each scale has been systematically sampled and included (✓), not systematically sampled but still included (∼), or not included entirely (×). All methods estimate functional diversity by combining species’ relative abundances in the community with the distributions of their trait values at the finest available scale. At that scale, the distribution of trait values of each species can be estimated directly (using a kernel density function) as a trait probability density function for that species (TPD_*S*_) if many conspecific individuals have been measured; if not, the TPD_*S*_ can be estimated as a multivariate normal distribution centred in the mean trait value, or the trait distribution can be substituted by the mean trait value entirely. Methods were applied to different taxa. For each taxon, the method using the highest available resolution in the trait data is indicated (*). See main text for full details of individual methods.

| Method | Name | Scale of trait variability included |  |  | Species' trait distributions are... | Resolution of trait data used | Taxa |
| --- | --- | --- | --- | --- | --- | --- | --- |
|  |  | Individuals | Populations | Species |  |  |  |
| M1 | TPD: individuals within populations | ✓ | ✓ | ✓ | estimated as TPD<sub>s</sub> directly from values | ≥10 values per population | Plants* |
| M2 | TPD: population mean | ✓ | ✓ | ✓ | estimated as TPD<sub>s</sub> centred in mean | ≥1 value per population | Plants |
| M3 | Classic: population mean | × | ✓ | ✓ | substituted by mean | ≥1 value per population | Plants |
| M4 | TPD: individuals across populations | ~ | ~ | ✓ | estimated as TPD<sub>s</sub> directly from values | ≥10 values per species | Plants, Ants* |
| M5a | TPD: species mean (sd for each species) | ~ | ~ | ✓ | estimated as TPD<sub>s</sub> centred in mean | ≥1 value per species | Plants, Ants |
| M5b | TPD: species mean (sd across species) | ~ | ~ | ✓ | estimated as TPD<sub>s</sub> centred in mean | ≥1 value per species | Ants |
| M6 | Classic: species mean | × | × | ✓ | substituted by mean | ≥1 value per species | Plants, Ants |

### Dimension reduction for ant traits and data preparation

We aimed to synthesize the major independent axes of variation in multidimensional trait space captured by the seven traits the ant dataset. The trait measurements (except body size) were first size-corrected by dividing by the measurement for body size (Weber’s Length). All traits were then log-transformed and standardized to have a mean of 0 and a standard deviation of 1. We performed a Principal Components Analysis (PCA) using the mean trait values of each species and subsequently predicted the values of the PCA components for all individuals in the dataset (after Martello et al., 2018). We retained the first two principal components, which had eigenvalues greater than unity, and predicted the values of these two components for each individual (N=319) in the trait dataset. The values for these two new ‘traits’ were used for all subsequent assessments of ant functional diversity. Trait data for plants were log-transformed and standardized prior to functional diversity calculations.

### Calculating functional diversity indices with different scales of trait variability

We used six different methods to calculate the FRic and Rao of every community (plot) in both the plant and ant datasets (Table 1). The methods differ generally in the scales of trait variability included in community functional diversity (Table 1), and specifically in the ways by which trait variability is scaled up to community functional diversity (see descriptions of individual methods below). They encompass a wide array of options available to researchers performing functional diversity analyses, and correspond to different scenarios, across which the required resolution of the trait data varies. The methods are ordered from high resolution (those systematically sampling trait values of individuals within populations and using these values to estimate probabilistic trait distributions of species directly) to low resolution (those using only the mean trait value of each species to calculate functional diversity indices).

### M1. TPD: individuals within populations

The various trait probability density (TPD) methods all involve generating a trait probability density function for each species – the TPD_*S*_ – which probabilistically summarises the distribution of a given species’ trait values at a given scale. The TPD_*S*_ functions of different species are then weighted using species’ relative abundances and aggregated to produce the trait probabilistic density function for the community at every plot – the TPD_*C*_. Functional diversity indices are then calculated based on properties of the TPD_*C*_ (for FRic) as well as the relationships between the TPD_*S*_ of different species (for Rao).

M1 includes the intraspecific trait variability among individuals in populations as well as among different populations. It can be used in a scenario where the traits of many individuals within each population have been measured systematically, such that one TPD_*S*_ for each population of a species can be estimated directly with a kernel density function and the trait values acquired from that population (Carmona et al., 2016a; 2019b).

To execute M1, we first estimated one TPD_*S*_ for each population of a species using kernel density functions and the trait values of conspecific individuals measured in each plot (henceforth we use ‘population’ in reference to conspecifics within the same plot). In doing so, intraspecific trait variability among individuals in the same population was captured by the TPD_*S*_ at a plot, while that among populations was captured by the TPD_*S*_ across different plots. We then aggregated the TPD_*S*_ of the different species in each plot (according to their relative abundances) to estimate the TPD_*C*_. We calculated FRic as the volume of the TPD_*C*_, and Rao from the dissimilarity among the TPD_*S*_ of the species present. A similar approach has been used in Carmona et al. (2019a).

### M2. TPD: population mean

This TPD method includes intraspecific trait variability among different populations. With a smaller degree of precision than M1, it also includes the intraspecific trait variability among individuals within the same population by likewise estimating one TPD_*S*_ for every population of a species. However, instead of using a kernel density function to estimate the TPD_*S*_ directly from the trait values of individuals (as in M1), M2 uses a variance estimation approach to estimate the TPD_*S*_ as a multivariate normal distribution centred in the mean trait value in the population (as proposed by Carmona et al., 2016a). It assigns the same among-individual trait variability to all species occurring within the same plot while doing this. This method is particularly relevant to a scenario where at least one individual from each population (plot) is measured, but the sample size is deemed insufficient for using a kernel density function to estimate TPD_*S*_ directly from the trait values acquired from the population (while there are no studies examining the minimum sample sizes for estimating TPD_*S*_ with different trait dimensions, Blonder 2016 recommends using at least a number of observations *m* such that log(*m*) > number of dimensions).

To execute M2, we first calculated the mean trait value in each population of each species, using the trait values of conspecific individuals measured in each plot. Then, we estimated the TPD_*S*_ using the *TPDsMean* function from the ‘TPD’ package (Carmona et al., 2019b) and the plug-in bandwidth (variance) selector implemented via the *Hpi*.*diag* function in the ‘ks’ package (Duong, 2015). The resultant TPD_*S*_ for each species in each plot was a multivariate normal distribution centred in the mean trait value of its local population, with a standard deviation determined by the estimated bandwidth (across all species in the same plot). Once the TPD_*S*_ were estimated, the aggregation of TPD_*S*_ to TPD_*C*_ at each plot, and the calculation of the FRic and Rao of each community were performed as in M1.

### M3. Classic: population mean

The various ‘Classic’ methods disregard the intraspecific trait variability among individuals in the same populations because they effectively assign the same trait value to all conspecific individuals in each plot. Classic methods calculate FRic using convex hulls and Rao using Gower-based dissimilarities. M3 includes intraspecific trait variability among different populations. It can be used in a scenario where at least one individual from each population (plot) is measured for each species (similar to M2).

To execute M3, we first calculated the mean trait value of each population of each species (as in M2). Next, we used those mean trait values of species in populations to calculate the FRic and Rao of the community at each plot directly (i.e., in contrast to TPD methods, which first estimated a TPD_*S*_ for each species and a TPD_*C*_ for each community). We calculated FRic using the convex hull method implemented in the dbFD function from the ‘FD’ package (Laliberté et al., 2014), and Rao with the Gower dissimilarity matrix between populations using the melodic R function (de Bello et al., 2016). A similar approach has been used in Carmona et al. (2015) and Gross et al. (2013).

### M4. TPD: individuals across populations

This TPD method includes intraspecific variability among the individuals of each species and treats the trait structure of species and communities in a probabilistic way, but does not strictly include the effects of differences between populations on traits. It can be used in a scenario where trait values have not been systematically acquired across the individuals and populations sampled, but the total sample size is nonetheless sufficient for using kernel density functions to directly estimate one TPD_*S*_ for the species as a whole (instead of one TPD_*S*_ per population, as in M1 and M2). For instance, some investigators may arbitrarily measure the traits of many individuals per species, or intentionally measure the traits of the smallest and largest individuals to capture the variability of the species, while ignoring the distribution of those individuals in the plots sampled.

To execute M4 we estimated one TPD_*S*_ for each species directly using a kernel density function and the trait values of all individuals measured across the different plots. At each plot, the aggregation of TPD_*S*_ to TPD_*C*_, and calculation of the FRic and Rao of the community were performed as in M1. A similar approach has been used in Traba et al. (2017).

### M5. TPD: species mean

This TPD method includes intraspecific trait variability of a similar structure as that of M4 but forgoes some precision in order to relax demands on sample size. Like M4, M5 estimates one TPD_*S*_ for each species only. However, it uses a variance estimator to estimate the TPD_*S*_ as a multivariate normal distribution centred in a mean trait value (like M2); in this case the overall species mean is used. M5 is thus particularly relevant to a scenario where insufficient individuals of each species have been measured to allow for a direct estimation of the TPD_*S*_ with a kernel density function.

To execute M5, we first calculated the mean trait value of each species from the trait values of all individuals measured across the different plots. We then used the *TPDsMean* function, as in M2 – but here we explored two alternative approaches for assigning the bandwidths (variances). These approaches reflect scenarios researchers encounter frequently.

In the first, **M5a. TPD: species mean (sd for each species)**, the bandwidth we used for estimating the TPD_*S*_ of each species was the standard deviation of all available trait values of that particular species (after Martello et al., 2018). This approach corresponds to the scenario where multiple trait values have been measured for each species, but the sample size is deemed insufficient for using a kernel density function to estimate TPD_*S*_ directly from those values. Still, this method assigns to each species an amount of intraspecific trait variability that reflects the trait differences between conspecifics observed in reality (Lamanna et al., 2014).

The second approach **M5b. TPD: species (sd across species)** was applied to the ant dataset only. This approach can be used in a scenario where only one trait value is available for each species – a common limitation in studies using trait information from the literature or databases. For this, we followed Lamanna et al. (2014) and estimated the TPD_*S*_ of every species using a constant bandwidth value: 0.5 times the standard deviation of the trait values across all species in the dataset. Once the TPD_*S*_ of all species were estimated via M5a or M5b, the aggregation of TPD_*S*_ to TPD_*C*_ at each plot, and the calculation of the FRic and Rao of each community were performed as in M1.

### M6. Classic: species mean

This method excludes intraspecific trait variability entirely and includes interspecific trait variability only. It is likely the most widely used approach in functional diversity assessments (Laliberté & Legendre, 2010), as it only requires a single trait value (e.g. the species-mean) for each species in the dataset and does not involve the estimation of trait probability density functions.

To execute M6, we first calculated the mean trait value of each species from the trait values of all individuals measured across the different plots. We then calculated the FRic and Rao of each community at each plot directly, following the same procedure as in M3.

All methods except M5b were applied to the plant data, while M4–M6 were applied to the ant data, because trait measurements of the ants were not associated with specific plots. We designated M1 and M4 as the HighRes models for plants and ants, respectively. All functional diversity analyses were performed in R (R Core Team, 2017); those involving TPD were performed using the ‘TPD’ package (Carmona et al. 2019b) while those involving convex hull volumes were performed using the ‘FD’ and ‘betapart’ (Baselga et al., 2018) packages.

### Statistical analyses

To investigate the relationships among different methods, we analysed the Pearson’s correlation between the values of each functional diversity index (FRic and Rao) calculated by the different methods with the plant and ant data. This allowed us to identify methods which yielded more similar results to the designated HighRes model overall.

We then investigated whether different methods captured the same ecological patterns. For the plant data, we investigated the changes in FRic and Rao in response to changes in water availability (%water content in soil samples taken from each plot) along the slope. For each index and method, we fitted a regression using water availability and its quadratic and cubic terms. For the ant data, we examined the changes in FRic and Rao in response to changes in percentage ground cover, and tested whether these patterns varied depending on the presence of the invasive species, *S. invicta*. For each index and method, we fitted a regression where we considered linear and quadratic terms for percentage ground cover, invasion status (binary variable: invaded/not invaded) and the interaction between them as predictors.

We used the ‘MuMIn’ package in R (Barton, 2016) to generate all potential subsets of all models, and ranked them using the Akaike information criterion corrected for small sample sizes (AICc). We selected the model from the HighRes method with the lowest AICc value as the one that best reflected the ecological patterns in each dataset. For each of the other candidate methods, we represented the results of the model with the lowest AICc value graphically, and calculated its ΔAICc (difference in AICc score) with respect to the selected HighRes model. An ΔAICc value of 0 indicates that the candidate method leads to a similar ecological interpretation as the HighRes method. Relatively small ΔAICc values (e.g. ΔAICc < 2) indicate that, while leading to different ecological interpretations, the HighRes model is not deemed as completely implausible under the candidate method. High ΔAICc values (e.g. ΔAICc > 2) imply that the functional diversity indices and their patterns produced by the candidate method lead to substantially different ecological interpretations from the results of the HighRes method (Burnham & Anderson, 2002).

## RESULTS

### Correlations between functional diversity indices calculated by different methods

In analyses for plants, the similarities between the HighRes method (M1) and the other methods (M2-M6) in their calculated FRic and Rao did not show a clear trend with the resolution of trait data and the scales of trait variability included (Fig. 1). Instead, for both indices, the values calculated with M4 were most similar to those calculated with M1 (ρ_Fric_ = 0.74; ρ_Rao_ = 0.83). All other methods largely failed to obtain FRic values similar to those from M1 (ρ ≤ 0.53) but performed better where Rao was concerned (ρ ≥ 0.73). In analyses for ants, the similarities between the HighRes method (M4) and the other methods (M5a, M5b, M6) in their FRic and Rao generally decreased with decreasing resolution in trait data and as fewer scales of trait variability were included (Fig. 2); M5a performed best, and especially well for Rao (ρ_Fric_ = 0.85; ρ_Rao_ = 0.99), while M6 performed the worst (ρ_Fric_ = 0.82; ρ_Rao_ = 0.66).

**Figure 1.**
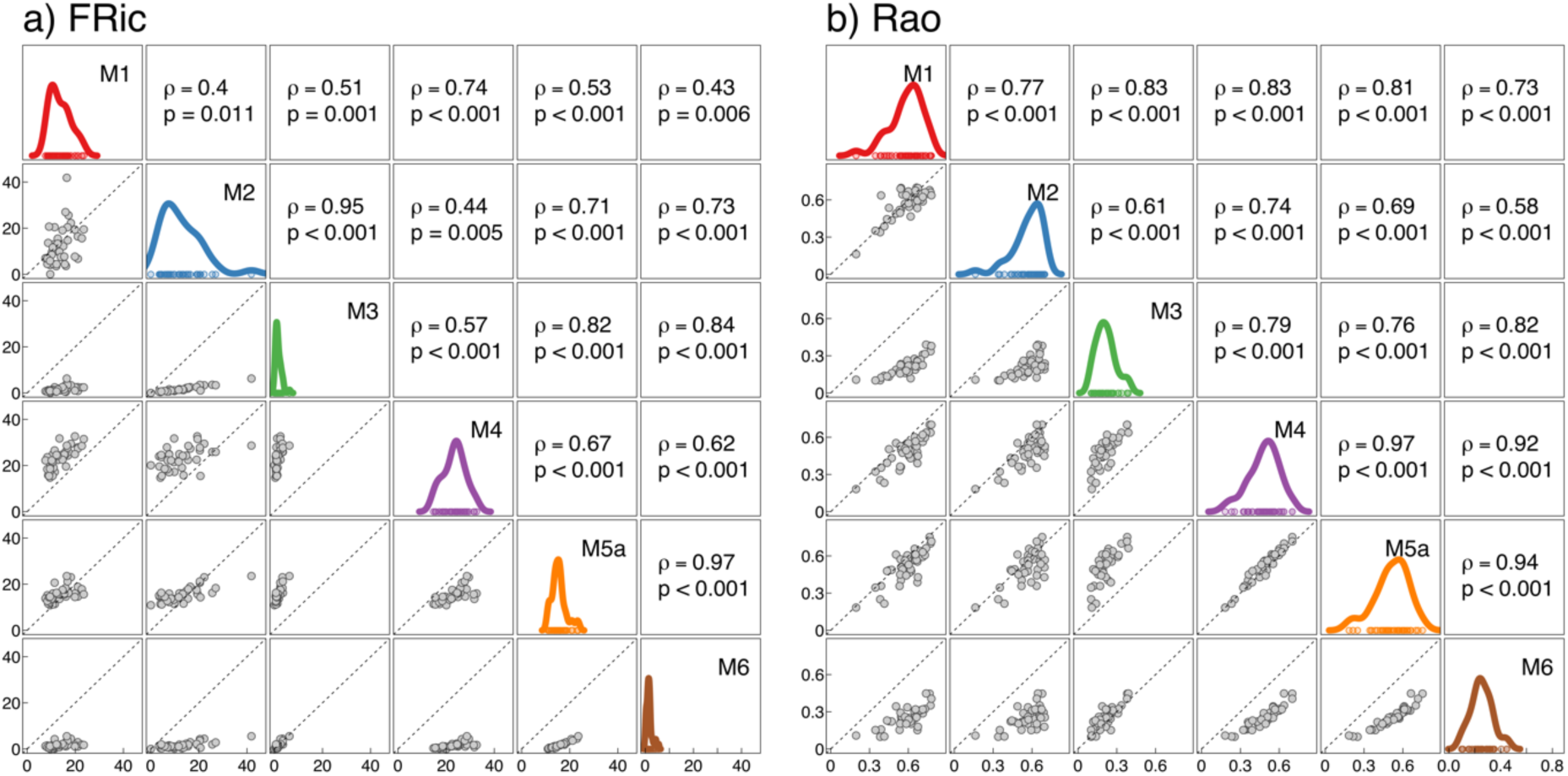
For 40 plant communities, plots show the degree of similarity, as measured by Pearson’s correlation (ρ), in values of Functional Richness (FRic) (a) and Rao (b) calculated by different methods. Each method includes a different scale (or scales) of trait variability in functional diversity, based on the resolution of the trait data (see Table 1). Plots along the diagonal depict probability density functions showing the distribution of FRic or Rao values calculated by individual methods.

**Figure 2.**
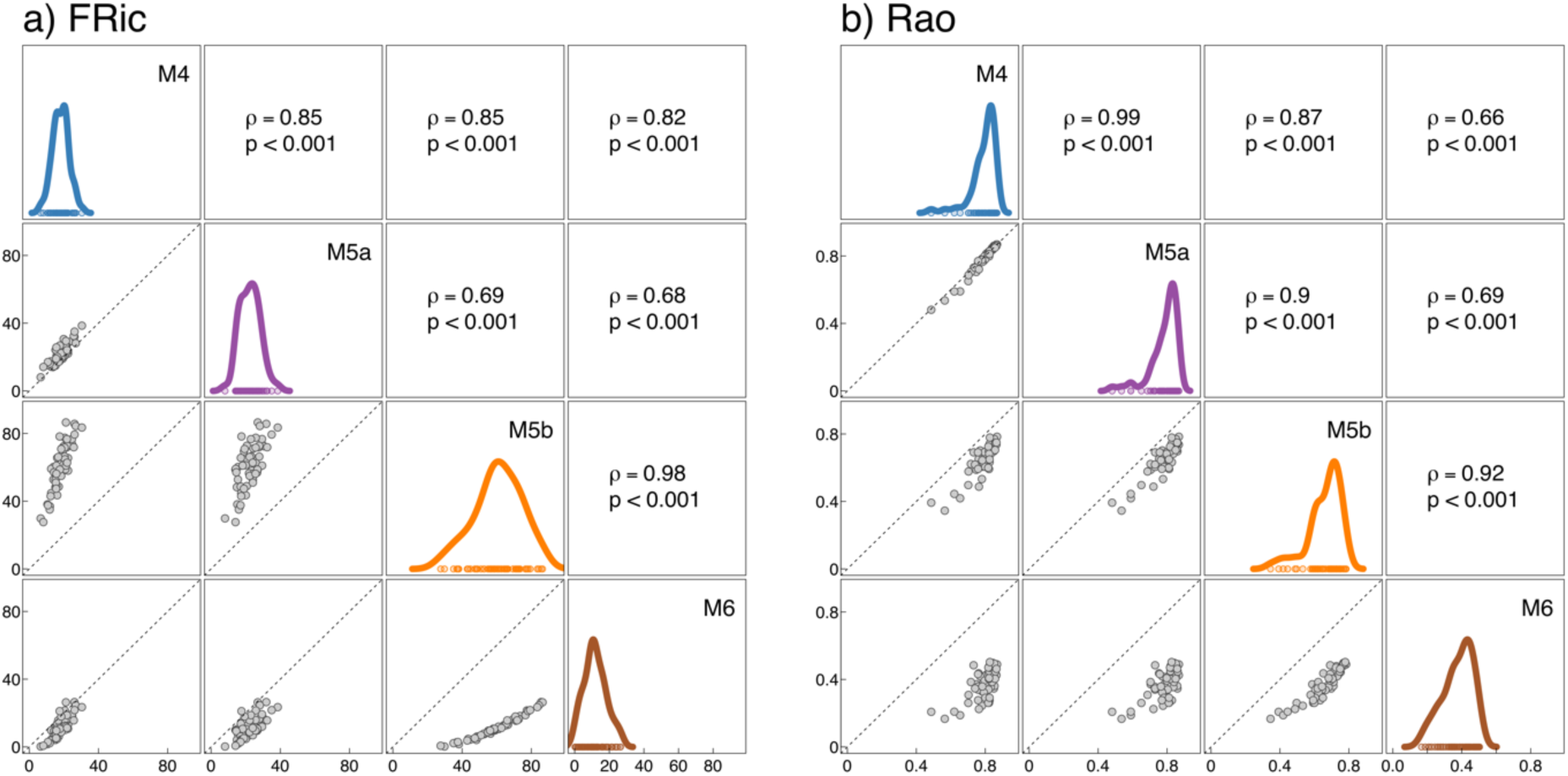
For 61 ant communities, plots show the degree of similarity, as measured by Pearson’s correlation (ρ), in values of Functional Richness (FRic) (a) and Rao (b) calculated by different methods. Each method includes a different scale (or scales) of trait variability in functional diversity, based on the resolution of the trait data (see Table 1). Plots along the diagonal depict probability density functions showing the distribution of FRic or Rao values calculated by individual methods.

### FRic responds to the exclusion of specific scales of intraspecific trait variability

The FRic patterns detected by the best models of the respective HighRes methods for both the plant and ant data were not likewise detected by the best models of the other methods which used trait data of lower resolution and which excluded particular scales of intraspecific trait variability. The plant model from the HighRes method (M1) detected a negative linear effect of soil water content on FRic (Fig. 3a). All other methods (M2-M6) led to substantially different ecological interpretations (Fig. 3b-f: ΔAICc > 2). The models from M3, M4 and M6 detected non-linear changes in which FRic peaked at intermediate soil water content (Fig. 3c,d,f), while those from M2 and M5a failed to detect an effect of soil water content on FRic (Fig. 3b,e). The ant model from the HighRes method (M4) detected a significant negative linear effect of ground cover on FRic and a significant effect of invasion (Fig. 4a). No models from the other methods detected an identical ecological pattern (Fig. 4b-d: ΔAICc ≠ 0). Though the model from M5b reproduced the significant negative linear effect of ground cover (Fig. 4c), it also detected a significant interaction effect between ground cover and invasion. The model from M6 failed to detect the effect of ground cover (Fig. 4d) while that from M5a failed to detect an effect of invasion (Fig. 4b).

**Figure 3.**
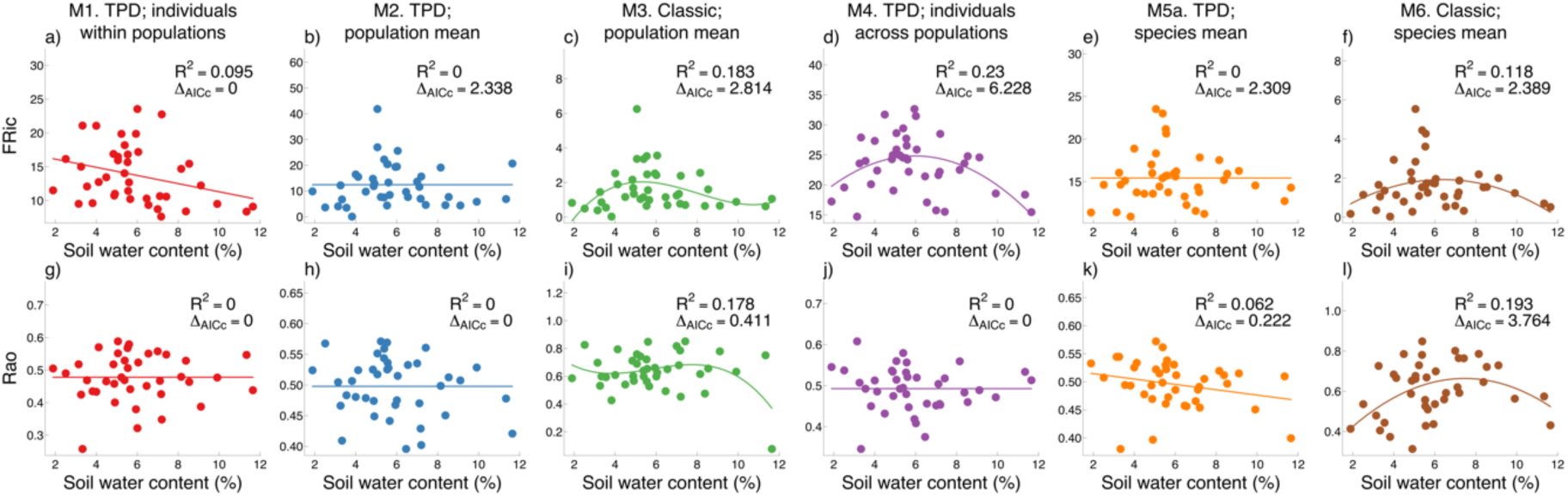
Six models of functional diversity measured in terms of the indices Functional Richness (FRic) (top) and Rao (bottom), in 40 Mediterranean plant communities (dots) distributed along a gradient of soil water content. The models were produced from six different methods for calculating functional diversity, each including a different scale (or scales) of trait variability, based on the resolution of the trait data available (see Table 1). For each index, the congruence of each model from M2-M6 with the model from M1 (which used data of the highest resolution) is summarised by an ΔAICc score, where a value of 0 indicates no distortion of the ecological pattern in the M1 model, and increasing values indicate increasing distortion.

**Figure 4.**
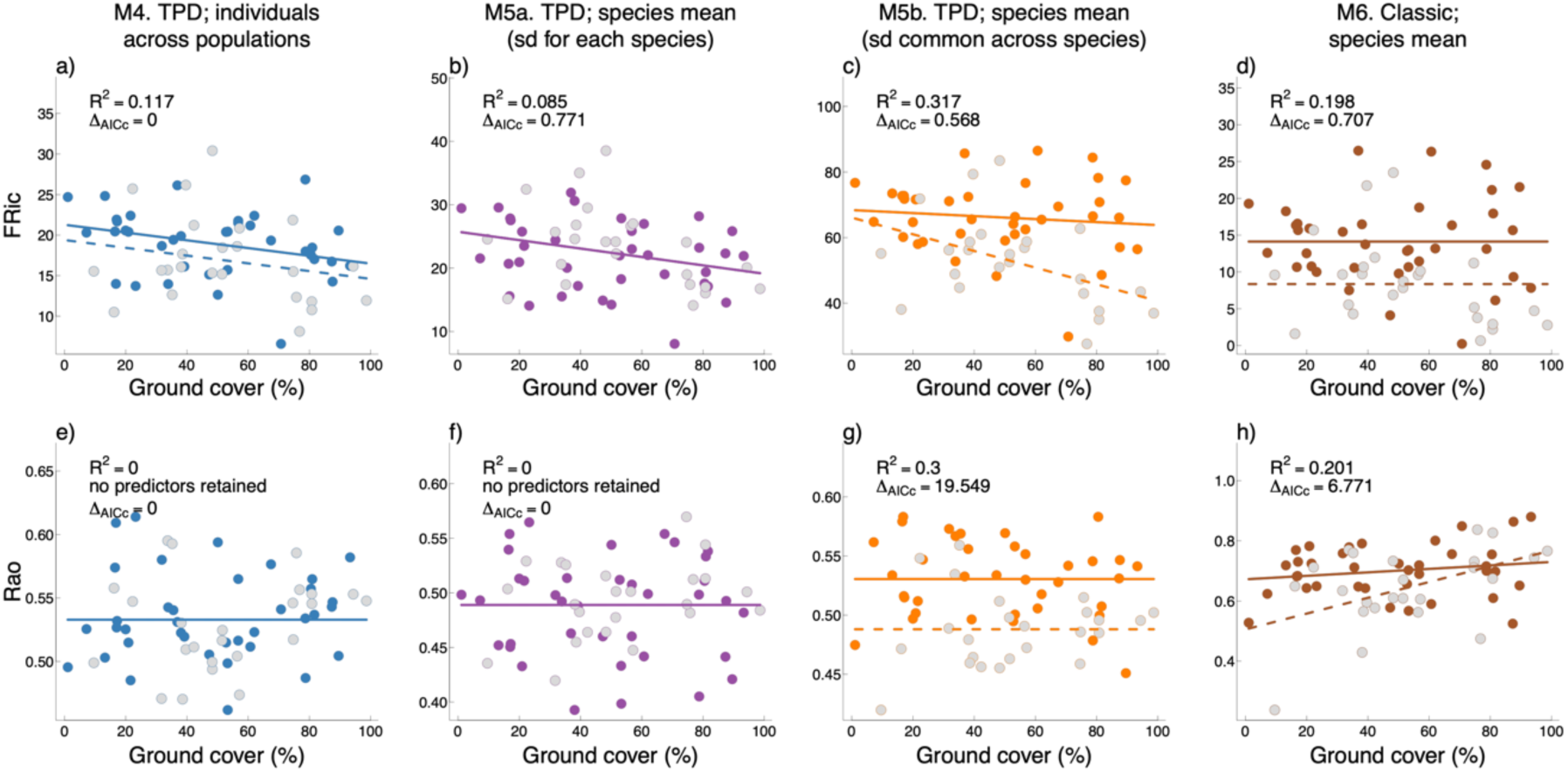
Four models of functional diversity measured in terms of the indices Functional Richness (FRic) (top) and Rao (bottom), in 61 ant communities distributed along a gradient of ground cover, including communities with (coloured dots, solid trend line) and without (grey dots, dotted trend line) an invasive species, *S. invicta*. The models were produced from four different methods for calculating functional diversity, each including a different scale (or scales) of trait variability, based on the resolution of the trait data available (see Table 1). For each index, the congruence of the models from M5a, M5b and M6 with the model from M4 (which used data of the highest resolution) is indicated by an ΔAICc score, where a value of 0 indicates no distortion of the ecological pattern in the M4 model, and increasing values indicate increasing distortion.

### Rao responds to the exclusion of intraspecific trait variability in general

The Rao patterns detected by the best models from HighRes methods were accurately reproduced by the best models from a few different methods which included intraspecific trait variability, but not by methods which excluded it entirely. The plant model from the HighRes method (M1) did not detect any significant effect of soil water content on Rao (Fig. 3g). This pattern was accurately reproduced by models from both M2 and M4 (Fig. 3h,j: ΔAICc = 0). In contrast, the model from M5a detected a significant negative linear effect of soil water content on Rao (Fig. 3k). Models from both M3 and M6 detected significant quadratic relationships between Rao and soil water content (Fig. 3i,l), with that from M6 leading to a substantially different ecological interpretation (Fig. 3l: ΔAICc > 2). The ant model from the HighRes method (M4) did not detect any significant effects from ground cover and invasion (Fig. 4e). This pattern was accurately reproduced with M5a only (Fig. 4f: ΔAICc = 0). By contrast, models from M5b and M6 detected significant effects with ground cover and invasion which led to very different ecological interpretations (Fig. 4g,h: 6 < ΔAICc < 20).

## DISCUSSION

Our empirical findings in two ecologically disparate systems show that widely used dissimilarity-based and volume-based indices capture different functional diversity patterns, and crucially, that these indices are strongly influenced by the particular scales of trait variability included, as determined by the specific resolution of the trait data available. These results imply that basic decisions in functional diversity assessments – those about whether traits are sampled *in situ*, and if so, from how many and which individuals; which indices are used and how they are calculated – can largely determine the patterns observed and even alter conclusions about ecological processes entirely. As our study examined the morphological traits of plants and ants, further work is needed to explore the effects of intraspecific trait variability on the functional diversity of other taxonomic assemblages and for traits spanning organisms’ physiology and behaviour. Nonetheless, in light of the findings, we suggest recommendations and issues to consider in both field-based and data-driven studies on functional diversity.

### Use multiple indices to draw inference

Limiting similarity (MacArthur & Levins, 1967) has long been invoked as a powerful driver of community structure, but the extent of its influence across taxonomic groups is less explored with trait-based approaches, which have the advantage of quantifying species’ niches in comparable terms (McGill et al., 2006). In our study, volume-based and dissimilarity-based functional diversity indices collectively detected patterns consistent with the effects of limiting similarity in the community structure of ecologically distinct groups such as plants and ants inhabiting different bioregions. Using trait data of the highest resolution, the best models in both groups showed that the total volume occupied by the trait values of all species (FRic) decreased along the respective environmental gradients, suggesting that the total niche space available to all members within the community was reduced (Fig. 3a, Fig. 4a). However, in spite of the shrinking niche space at the community level, the overlap between species within this space – as measured by their trait dissimilarity, Rao – remained constant (Fig. 3g, Fig. 4e). The limiting similarity hypothesis is further supported by evidence that species richness declined along the environmental gradient in both cases (Supporting information). To the best of our knowledge, this is the first study to show the importance of including intraspecific variability at the within-plot scale for detecting the effects of limiting similarity in natural assemblages. It supports previous findings in simulated (de Bello et al., 2013) and experimental conditions (Mason et al., 2011). More broadly, these results demonstrate that using different indices to target distinct facets of functional diversity can enhance inferences about ecological processes. Nonetheless, the relationships were detected using trait data of high resolution, which is not always available. Moreover, we found functional diversity indices to be very sensitive to the particular scales of trait variability included. An understanding of these relationships is therefore crucial for selecting appropriate methods and avoiding misinterpretation in functional diversity research.

### Results from mean-based methods may distort patterns and alter inference

In keeping with previous studies (Baraloto et al., 2010; de Bello et al., 2013), we found that calculations solely using the mean trait values of species misestimated functional diversity (Fig. 1: values from M1 are weakly correlated with those from M6). Our results also go further than previous work, as they empirically show that mean-based methods for calculating functional diversity *in general* – such as convex hull-based FRic and Gower-based Rao – can distort patterns and alter conclusions about underlying processes. For instance, a negative-linear relationship may be transformed to a quadratic one (Fig. 3a vs. Fig. 3f), a ‘false’ effect of an invasive species may be detected (Type I error) (Fig. 4e vs. Fig. 4h), and a ‘true’ effect of an environmental gradient may fail to be detected (Type II error) (Fig. 4a vs. Fig. 4d). Furthermore, even when an effort is made to sample among-population trait variability, such as by systematically sampling the traits of conspecific individuals within plots, the resultant functional diversity patterns may still be distorted if the among-population trait variability is derived solely from the mean trait values of populations (Fig. 3: in both FRic and Rao, the model from M3 clearly changes the relationship modelled with M1). Given these apparent limitations of mean-based methods in preserving the integrity of functional diversity estimates and patterns, calls for using probabilistic methods to include intraspecific trait variability into functional diversity (Carmona et al., 2016a; Blonder et al., 2018) should not be understated. Yet, our results show that both the scales of intraspecific trait variability considered (among individuals vs. among populations) and the precise methods with which probabilistic trait distributions are estimated (e.g. M1, M2, M4, M5a, M5b) will strongly influence the functional diversity observed.

### Aim to estimate probabilistic trait distributions from trait values of individuals directly

We recommend that ecologists sample traits of multiple individuals per species and use these values to estimate probabilistic trait distributions for each species directly with kernel density functions (e.g. in M4), even when this cannot be done for every population of every species (as in M1). In our study, such an approach sampling at least 10 individuals of each species (M4) achieved values of FRic and Rao that were most similar to those calculated by the method using trait data of the highest resolution, which sampled at least 10 individuals per population of each species (M1) (Fig. 1).

Our results show that FRic is especially sensitive to trait variability among individuals within the same populations, and we suggest that assessments of FRic and community niche space can address this scale of trait variability by sampling the traits of conspecific individuals occurring within the same plots. Here, the FRic of plant communities was overestimated by methods which either estimated the trait probability density functions of species (TPD_*S*_) from mean values or methods which used only the mean value at the population or species levels (M2, M3 and M6), as compared to methods which estimated TPD_*S*_ using the trait values of individuals directly (M4) (Fig. 1a: compare plots for M1 against M2, M3 and M6, with plots for M1 against M4). These relationships imply that each plant community actually occupied a smaller niche space than would be expected if it only included the among-population trait variability in each species, or if it excluded intraspecific trait variability entirely. This reduction in community niche space locally may be driven by environmental filtering or competition hierarchies promoting similarity in the traits of coexisting individuals (Germain et al., 2018; Carmona et al., 2019a). Yet our findings demonstrate that such important assembly processes may fail to be detected if among-individual trait variability is not included in functional diversity assessments.

The low correlation between the FRic values of M1 and M2 (Fig. 1a) was particularly surprising. Both methods capture trait variability at the same spatial and organizational scales (Table 1); thus, the low correlation arose from the different ways by which the TPD_*S*_ of populations were estimated. That estimating TPD_*S*_ directly using kernel density functions and all sampled values (M1) and estimating them as multivariate normal distributions centred in the mean sampled values (M2) produced distinct results shows that FRic is strongly influenced by the effects of individuals with extreme trait values (in spite of TPD methods are theoretically more robust to such effects than convex hull-based hypervolumes; Carmona et al., 2016a). This sensitivity of FRic to extreme values could be investigated further by testing the effects of setting smaller probability thresholds to the TPD_*S*_ (see Blonder, 2016 and Carmona et al. 2016b).

We found that Rao, like FRic, was sensitive to the exclusion of intraspecific trait variability overall (Fig. 1b: M6 vs. M1; Fig. 2b: M6 vs. M4; Fig. 3g vs. 3l; Fig 4e vs. Fig. 4h). Yet, in contrast to the patterns observed for FRic, the methods estimating species’ trait probability density functions directly from individual values using kernel density functions (M1, M4) and those estimating them as multivariate normal distributions centred in the mean trait value (M2, M5a) produced similar levels of Rao (Fig. 1b: M1 vs. M2 for plants; Fig. 2b: M4 vs. M5a for ants). Furthermore, Rao calculated by these different methods followed similar trends with environmental gradients (Fig. 3g,h,j and Fig. 4e,f). These results are encouraging – although distinct, the two methods are evidently valid alternatives for calculating trait dissimilarities between populations or species. The results also suggest that estimations of dissimilarities between species (e.g. with Rao) are more robust to methodological choices than estimations of the total functional space occupied by communities (e.g. FRic).

Observed functional diversity will most likely approximate reality when TPD_*S*_ are estimated directly using the trait values of multiple individuals per species and information on the spatial structure of their populations across environments (Carmona et al., 2016a). Yet, large samples of trait measurements and spatially-detailed trait information are seldom available. With respect to these limitations of empirical studies, our results suggest that even with a lower resolution in trait data, intraspecific trait variability can still be included in dissimilarity-based functional diversity without leading to significant distortion in ecological patterns. This is in agreement with previous results suggesting that trait sampling is more efficient when local (i.e., measured in the corresponding plot) trait values are used, rather than a regional average for each species (Carmona et al., 2015, Gross et al., 2013, Baraloto et al., 2010). In the specific case of the TPD framework, this can be achieved by using variance estimators to estimate the TPD_*S*_ of each species as a multivariate normal trait distribution centred in the sampled mean trait value (e.g., M2, M5a).

In estimating the TPD_*S*_ of species with variance estimators, however, one should not assume that the same algorithm is valid for estimating the variance in the distributions of trait values of disparate taxa. This is evidenced by our finding that the FRic of ant communities from when TPD_*S*_ was estimated assuming a constant variance (following the solution proposed in a plant-based study; Lamanna et al., 2014) (M5b) was over two times higher than their FRic from when TPD_*S*_ was estimated using the trait values of all individuals observed (M4), as well as that from when TPD_*S*_ was estimated using the variance in the data (M5a) (Fig. 4a-c). Specifically, this suggests that in comparison to plants, trait variation within ant species may be very low relative to trait variation between species (as reported recently in Gaudard, Robertson & Bishop, 2019). In any case, these results strongly suggest that decisions about the precise variances used for estimating trait distributions should be grounded in the ecological characteristics of the studied organisms. Caution is therefore advised for multi-taxa analyses, where it may seem temptingly efficient to apply the same estimation procedure across all groups. Further studies on the contribution of intraspecific variability to total trait variability in different taxa (e.g. Siefert et al., 2015) will be useful for navigating these issues.

## Conclusion

This study demonstrates the strong influence that trait variability within species can have on our view of functional diversity and underlying ecological processes. It also clarifies when and how intraspecific trait variability can be reasonably included in functional diversity studies with limited trait data. Accounting for this crucial yet overlooked source of functional variability in nature will be key to understanding the responses and effects of biodiversity.

## ACKNOWLEDGEMENTS

MKLW is supported by a grant from the National Geographic Society (60-16) and a Clarendon Scholarship from the University of Oxford. CPC was supported by the Estonian Research Council (project PSG293) and the European Union through the European Regional Development Fund (Centre of Excellence EcolChange). The authors thank Owen Lewis and Benoit Guénard for comments on the manuscript.

